# Phytohormone dynamics during dormancy progression in apricot (*Prunus armeniaca* L.): responses to chilling deficiency

**DOI:** 10.64898/2026.11.27.769669

**Authors:** Alvaro Delgado, Hisayo Yamane, Raquel Sánchez Pérez, Tzu-Fan Hsiang, Mikiko Kojima, Hitoshi Sakakibara, Juan Alfonso Salazar, Jose A. Egea, David Ruiz

**Author notes:** Corresponding author: Alvaro Delgado.

## Abstract

Adequate fulfilment of the chilling requirement (CR) is essential for synchronized flowering and stable fruit production in temperate fruit trees, but hormonal responses to insufficient chilling remain poorly understood. We characterized phytohormone dynamics in apricot (*Prunus armeniaca* L.) floral buds through two experiments over consecutive seasons: three cultivars with contrasting CRs evaluated at warm and mild/cold sites, and a high-chill cultivar exposed to ambient conditions or warming-induced chill deprivation within one orchard. Profiling of up to 25 phytohormones around CR fulfilment revealed broadly conserved temporal trajectories, with differences in response magnitude and timing. Abscisic acid (ABA) declined under both adequate and insufficient chilling, indicating that its decline alone may not distinguish successful dormancy progression from impaired development. Gibberellin A₁ (GA₁) remained low before CR fulfilment and increased during subsequent growth. Increases in selected cytokinins, jasmonic acid (JA), auxins, and GA₁ accompanied growth resumption under adequate chilling. N⁶-(Δ²-isopentenyl)adenine (iP) showed the most reproducible increase around or after endodormancy release across cultivars and experiments, supporting its potential as a candidate indicator. Across experiments, insufficient chilling produced consistent mean decreases in JA, trans-zeatin (tZ), and dihydrozeatin riboside 5J-monophosphate (DZRMP), and increases in cis-zeatin O-glucoside (cZOG) and trans-zeatin-9-N-glucoside (tZ9G). Although growth partially resumed under chill-deficient conditions, several hormonal responses were attenuated or delayed, particularly under severe deprivation, alongside previously reported reductions in bud viability and cumulative reproductive success in the same trees. These findings identify candidate hormonal indicators of endodormancy release and chill deficiency, requiring validation across broader genetic and environmental conditions before practical application.

## 1. Introduction

Bud dormancy is a fundamental adaptive mechanism that enables temperate deciduous fruit trees to survive adverse winter conditions and synchronize vegetative and reproductive growth in spring (Faust et al., 1997; Saure, 1985). This process is traditionally divided into two phases: endodormancy, during which bud growth inhibition is regulated by internal factors within the flower and vegetative buds, and ecodormancy, during which growth is constrained by external environmental conditions (Lang et al., 1987). During endodormancy, buds require exposure to prolonged periods of low temperature to satisfy their cultivar-specific chilling requirement (CR), whereas during ecodormancy, growth resumption depends mainly on the accumulation of sufficient heat (i.e., heat requirements; HR), with both phases often overlapping depending mainly on the developmental stage and environmental conditions (Delgado et al., 2025b; Pope et al., 2014). In temperate fruit crops, adequate fulfilment of the genotype-specific CR is essential for successful endodormancy release, normal flower development, and stable fruit production (Atkinson et al., 2013; Campoy et al., 2011; Delgado et al., 2026a). Among these species, apricot (*Prunus armeniaca* L.), like most fruit crops of the genus *Prunus*, is particularly sensitive to insufficient winter chill. Because the satisfaction of cultivar-specific CR is the main determinant of climatic adaptation (Delgado et al., 2025a; Luedeling, 2012), the ongoing increase in winter temperatures represents a significant threat to the long-term sustainability of fruit production in traditional growing regions (Fernandez et al., 2023). Therefore, accurate identification of endodormancy release and the subsequent determination of the corresponding CR values is essential for evaluating cultivar adaptation and predicting orchard performance under changing climatic conditions. Nevertheless, its empirical identification remains challenging because dormant buds show limited external morphological changes during this period (Dennis, 2003; Fadón and Rodrigo, 2018). As a result, substantial research efforts have been made to identify physiological, metabolic, and molecular markers associated with dormancy progression and release in temperate fruit species, including almond, peach, apricot, sweet cherry, pear, and apple (Del Cueto et al., 2017; Fadón et al., 2023; Fernandez et al., 2019; Guillamón et al., 2020; Ionescu et al., 2017; Yu et al., 2020). Despite substantial advances in understanding bud dormancy, the regulatory molecular and physiological mechanisms controlling its induction, maintenance, and release remain only partially elucidated (Yamane et al., 2023). Among the physiological regulators involved in plant dormancy, phytohormones play a central role by integrating environmental cues, such as temperature, with developmental responses (Wilkie et al. 2008). A diverse array of phytohormones interact dynamically to coordinate growth, development, and adaptive responses throughout the plant life cycle (Davies, 2004). In fruit trees, these interactions also regulate flower bud differentiation and reproductive development (Wilkie et al. 2008). Current evidence indicates that dormancy progression and bud burst are controlled by complex interactions among hormonal signalling, carbohydrate metabolism, and redox homeostasis (Considine and Foyer, 2014). Carbohydrates provide essential carbon sources and energy reserves for renewed bud growth, with starch progressively degraded into soluble sugars during dormancy progression (Fadón et al., 2018; Hernandez et al., 2021; Ito et al., 2012). In parallel, alterations in oxidation–reduction processes have been associated with the transition from dormant to active growth (Takemura et al., 2015).

In this sense, advances in metabolomic and molecular approaches have provided valuable insights into these regulatory networks and facilitated the identification of biomarkers associated with endodormancy release. For example, in almond, metabolomic analyses have identified compounds such as ascorbic acid and prunasin as potential indicators of endodormancy release (Guillamón et al., 2020), while the progressive downregulation of *DORMANCY-ASSOCIATED MADS-box* (*DAM*) genes has consistently been associated with the transition from endodormancy to active growth in several *Prunus* species (Delgado et al., 2026b; Falavigna et al., 2019; Quesada-Traver et al., 2020).

Among phytohormones, research on dormancy release and flowering in fruit trees has primarily focused on abscisic acid (ABA) and gibberellins (GAs) as they constitute the central regulatory axis controlling dormancy maintenance and release in woody perennials (Cooke et al., 2012; Hernandez et al., 2021; Howe et al., 2015; Zheng et al., 2015). Although the role of GAs remains controversial, with both promotive and inhibitory effects on bud break being reported (Ionescu et al., 2017; Zheng et al., 2018), a progressive decline in ABA levels appears to be a prerequisite for endodormancy release and has been proposed as a reliable indicator in species such as *Pyrus pyrifolia*, *Prunus persica*, *Prunus armeniaca* and *Populus* spp. (Guillamón et al., 2025; Howe et al., 2015; Ito et al., 2021; Yan et al., 2025). Furthermore, increasing evidence suggests that the ABA/GA balance, rather than the absolute concentration of either hormone, is a key determinant of dormancy status in species such as sweet cherry and Japanese apricot (Liu and Sherif, 2019; Zheng et al., 2018). Auxin signalling has also been implicated in chilling-mediated developmental transitions, although its precise role in dormancy regulation has received less attention than ABA-related pathways (Beauvieux et al., 2018).

On the other hand, cytokinins have emerged as overlooked regulators of dormancy release in fruit tree species despite being essential regulators of meristem activation, cell division and shoot development (Sakakibara, 2006). Their contribution to dormancy regulation, particularly under conditions of insufficient chilling, remains poorly understood, and available evidence is often contradictory. In sweet cherry, no major differences were observed in cytokinin bases or conjugated forms between endodormancy and ecodormancy phases (Götz et al., 2023), whereas hydrogen cyanamide treatments induced significant increases in specific cytokinin metabolites prior to bud burst (Bound et al., 2022; Ionescu et al., 2017). In apple, cytokinin signalling has been shown to interact with flowering regulators such as SOC1-like genes, suggesting a role in developmental reactivation after dormancy release (Li et al., 2019). However, interpretation of these findings is further complicated by the diversity of cytokinin forms, including free bases, ribosides, precursors, and conjugates, which are rarely quantified comprehensively (Romanov and Schmülling, 2022). Consequently, whether cytokinins act as key integrators of the transition from dormancy to active growth remains unresolved.

Furthermore, proposed physiological and molecular markers are still rarely incorporated into breeding programmes or cultivar evaluation strategies (Yang et al., 2021). This knowledge gap is particularly relevant under climate change scenarios, as insufficient winter chilling can disrupt bud development, reduce flowering intensity, and impair fruit set, while the molecular and hormonal processes driving these responses remain poorly characterized. Recent evidence in apricot indicates that *FLOWERING LOCUS T (FT),* rather than *DAM* genes, may provide a more reliable indicator of the effects of suboptimal chilling on flowering performance (Delgado et al., 2026b), suggesting that chilling deficiency may induce specific molecular responses distinct from those associated with dormancy release under optimal conditions. Likewise, little is known about how reduced chilling modifies hormonal dynamics, including the balance between cytokinins and acidic phytohormones, throughout dormancy progression. Current knowledge is further limited because many studies investigating hormonal regulation of dormancy have been conducted under controlled environmental conditions using young plants (Borkowska and Powell, 1982; Endo et al., 2018; Ito et al., 2021), whereas information from mature, productive trees exposed to natural agroclimatic variability remains scarce. Consequently, the extent to which chilling deficits alter endogenous hormonal networks and contribute to flowering disorders such as reduced floral density, bud abscission and poor fruit set, remains largely unknown.

To address these knowledge gaps, in this work we examined phytohormone dynamics in apricot floral buds through two complementary experiments conducted over consecutive seasons in apricot (*Prunus armeniaca* L.). Three cultivars with contrasting chilling requirements were first evaluated at warm and mild/cold sites, followed by a semi-controlled chill-deprivation experiment involving a high-chill cultivar. Apricot represents a relevant model due to its high sensitivity to insufficient winter chill and its relatively limited study compared with other fruit tree species (Delgado et al., 2026c, 2025a). Particular emphasis was placed on cytokinins by quantifying their bioactive forms, precursors, and catabolites throughout dormancy progression. Simultaneously, endogenous levels of several acidic phytohormones were also monitored to provide a comparative framework for interpreting cytokinin dynamics during seasonal dormancy transitions.

By examining multi-hormonal profiles in the context of previously reported phenological and reproductive responses to contrasting chilling conditions, this study aimed to (i) determine whether temporal changes in phytohormone concentrations during the late stages of dormancy can identify the timing of endodormancy release, and (ii) assess how chilling deficits alter hormonal networks and may contribute to flowering disorders in apricot. Overall, this study provides insights into phytohormone dynamics during dormancy progression and supports the identification of potential physiological indicators of chilling fulfilment in apricot.

## 2. Materials and Methods

### 2.1. Plant material, experimental design, and agroclimatic conditions

The study comprised two complementary experiments conducted over two consecutive dormancy seasons (2023/2024 and 2024/2025) using adult, fully productive apricot (*Prunus armeniaca* L.) trees. During the first season, the plant material included three self-compatible cultivars spanning contrasting CRs and flowering dates: ‘Cebasred’ (low-chill), ‘Búlida’ (medium-chill), and ‘Flopría’ (high-chill). All three cultivars were monitored simultaneously at two climatically contrasting locations in the Murcia Region (southeastern Spain) with differing levels of chill accumulation: Archena (38°05J49.6JN, 1°19J30.2JW; 145 m a.s.l.) and Cieza (38°16J13.3JN, 1°34J01.6JW; 250 m a.s.l.), previously characterized as warm and mild/cold environments, respectively (Delgado et al. 2025a). During the second season, a semi-controlled chill-deprivation experiment was conducted at the same mild/cold site to test whether the hormonal patterns observed during the first season were reproducible while minimizing confounding effects associated with differences between orchards, such as soil conditions, tree age and orchard management. For this experiment, we selected the high-chill, self-compatible cultivar ‘Ladycot’ due to its comparable CR to ‘Flopría’. Based on the Dynamic Model (Fishman et al., 1987a, 1987b), ‘Ladycot’ and ‘Flopría’ had chilling requirements of 57 and 58 Chill Portions (CP), respectively, under the conditions of the present study (Delgado et al., 2026a, 2025a). The chill-deprivation experiment consisted of: (i) chill-deprived trees subjected to long-term warming under a plastic-film cover and (ii) trees exposed to ambient winter conditions (uncovered control). A detailed description of the plastic-film covering system is provided by Delgado et al. (2026a). Hourly air temperature was recorded throughout both experimental seasons. During the first season, temperature data were obtained from the nearest weather stations belonging to the Murcia Agrarian Information System (SIAM-IMIDA). During the second season, Testo 174 H mini data loggers were installed inside and outside the plastic-film structure. To ensure comparability between seasons, logger measurements were adjusted against concurrent records from the official SIAM-IMIDA weather station at the experimental site. Chill accumulation was calculated from the standardized hourly temperature data recorded between 1 November and 15 March using the Dynamic Model (Fishman et al., 1987a, 1987b) and expressed as CP.

### 2.2. Determination of endodormancy release and floral bud sampling

Endodormancy release was determined using the bud-forcing method described by Ruiz et al. (2007) and the chill accumulated up to this date was used to estimate cultivar-specific CRs. Floral buds were sampled from multiple positions within a single mature tree to minimize potential effects of bud orientation. At each sampling date, buds were pooled into three independently processed samples to capture within-tree phenological variability.

During the first season, sampling was scheduled at late phases of endodormancy and the transition towards ecodormancy. Because the cultivars differed in their CRs, the number of sampling dates was three dates for ‘Cebasred’, four for ‘Búlida’, and five for ‘Flopría’ (Table S1). Except for ‘Cebasred’, buds were collected from visually dormant stages (Baggiolini stage B) through early green tip (stage B–C; Baggiolini, 1952). ‘Cebasred’ was sampled only after endodormancy release, at approximately 105, 115, and 130% CR fulfilment. ‘Búlida’ was sampled during late endodormancy and its release (80–100% CR fulfilment) and subsequent development (100–113%). Sampling in ‘Flopría’ extended from approximately 67 to 108% CR fulfilment, covering a broader portion of the dormancy transition than in other cultivars. As chill accumulation and dormancy progression increasingly diverged between locations, its final three samplings at the warm site were delayed by one week to improve stage comparability with the mild/cold site. ‘Cebasred’ and ‘Búlida’ were sampled on the same dates at both locations.

During the second season, the semi-controlled chill-deprivation experiment began sampling earlier to capture hormonal dynamics during deeper endodormancy. The four sampling dates were initially scheduled to represent approximately 60, 90, 105, and 120% of the historical CR of ‘Ladycot’, estimated over three previous seasons at the same location (Delgado et al., 2026a). Because interannual variation in temperature influences chill accumulation and dormancy progression, CR fulfilment was subsequently recalculated using the endodormancy release date observed during the experimental season. The sampling dates thus corresponded to approximately 63, 94, 110, and 126% of the season-specific CR in control trees (Table S2). These sampling points covered developmental stages broadly comparable to those examined during the first season, particularly in the high-chill cultivar. The plastic-film treatment prevented the chill-deprived trees from satisfying their cultivar-specific CR. Consequently, control and treated trees were sampled on the same calendar date (Table S2). In both seasons, collected buds were immediately placed in Eppendorf tubes, frozen on dry ice in the field, transferred to liquid nitrogen, and stored at −80 °C in the laboratory until further analysis.

### 2.3. Phytohormone extraction and quantification

Flower bud samples were freeze-dried for 72 h using a LyoBeta 25™ freeze-dryer (Telstar, Spain) and they were subsequently stored at −20 °C in the dark until analysis. Lyophilized floral buds were ground to a fine powder using a multi-beads shocker (Yasui Kikai Co., Osaka, Japan). Phytohormones were extracted, purified and separated into three fractions as previously described (Kojima et al., 2009; Kojima and Sakakibara, 2012). Automated solid phase extraction (SPE) was performed using either a Hamilton STAR robotic workstation (Hamilton, Reno, NV, USA) or a Fluent 780 workstation (Tecan, Männedorf, Switzerland), equipped with Oasis HLB, MCX and MAX 96-well SPE plates (30 mg; Waters, Milford, MA, USA). The resulting three fractions were used for the analysis of acidic phytohormones, cytokinin mononucleotides, and cytokinin bases, ribosides, and glucosides respectively. Chromatographic separation and mass spectrometric detection were performed using an ACQUITY Premier UPLC system coupled to a Xevo TQ-XS triple quadrupole mass spectrometer (Waters). Cytokinin nucleotides were separated on an Ascentis Express F5 column (150 × 3 mm, 2.7 µm; Merck, Darmstadt, Germany) using 0.1% formic acid in water (solvent A) and acetonitrile (solvent B) as the mobile phases. Other cytokinin species, including free bases, ribosides, and glucosides, were separated on an ACQUITY Premier HSS T3 VanGuard FIT (2.1 × 100 mm, 1.8 µm; Waters) using 0.06% acetic acid in water (solvent A) and 0.06% acetic acid in 70% methanol (solvent B) as the mobile phases. Acidic phytohormones, including gibberellins (GAs), abscisic acid (ABA), salicylic acid (SA), jasmonic acid (JA), jasmonoyl-L-isoleucine (JA-Ile), indole-3-acetic acid (IAA), and auxin conjugates, were separated on an ACQUITY Premier HSS T3 VanGuard FIT (2.1 × 100 mm, 1.8 µm; Waters) using 0.1% formic acid in water (solvent A) and acetonitrile (solvent B) as the mobile phases. Data were processed using TargetLynx software within MassLynx version 4.2 (Waters). Phytohormone concentrations were expressed as pmol g⁻¹ dry weight (DW).

The same analytical panel was used in both seasons, although the compounds that could be reliably quantified differed. During the first season, these primarily comprised cytokinins, together with indole-3-acetyl aspartate (IAA-Asp), JA, and SA. During the second season, cis-zeatin riboside 5J-monophosphate (cZRMP) and trans-zeatin-7-N-glucoside (tZ7G) could not be reliably quantified, whereas dihydrozeatin-9-N-glucoside (DZ9G), ABA, gibberellin A₁ (GA₁), IAA, and JA-Ile were successfully quantified in addition to the compounds quantified in both seasons. The ABA/GA₁ ratio was subsequently calculated from their quantified concentrations. The complete list of phytohormones and their quantification status in each experimental season is provided in Table S3.

### 2.4. Agroclimatic, phenological, and reproductive performance data

Agroclimatic, phenological, and reproductive data from the same trees and experimental conditions were previously reported by Delgado et al. (2026b, 2026a) and were included here to provide context for interpreting the phytohormone profiles of the two high-chill cultivars that exhibited flowering disorders. The selected variables comprised chill accumulation in the experimental period, full bloom date, viable floral buds, pistil abortion, fruit set and cumulative reproductive success, calculated as the number of small fruits relative to the initial number of dormant floral buds. Detailed assessment procedures are provided in the original studies (Delgado et al., 2026b, 2026a).

### 2.5. Statistical analysis

Three independently processed bud pools from a single tree were analysed per sampling date and experimental condition. In the first season, one replicate was shipped separately and showed recurrent discrepancies potentially associated with transport conditions. Potential outliers were assessed separately for each phytohormone by visual inspection of replicate values. At most one observation per phytohormone, sampling date, and condition was excluded when it differed markedly from the other two replicates. Principal component analysis (PCA) was performed on mean phytohormone concentrations for each cultivar × agroclimatic area × sampling-point combination using the FactoMineR package in R. For heatmap visualization, mean concentrations were standardized separately for each phytohormone across all cultivar, agroclimatic area, and sampling-point combinations. The resulting Z-scores indicate the number of standard deviations by which each value deviates above or below the overall mean for that phytohormone.

The two experimental seasons were analysed separately, and ANOVA was performed only for the second season. For each phytohormone, log₁₀(x + 1)-transformed concentrations were analysed using Type III ANOVA with treatment, sampling date, and their interaction as fixed effects. Residual normality and variance homogeneity were assessed using Shapiro–Wilk and Levene’s tests, respectively. Estimated marginal means were compared between treatments within each date and between dates within each treatment using Tukey-adjusted P-values.

Exploratory Spearman correlations used mean concentrations for each season × cultivar × sampling-point combination from the first-season mild/cold site and second-season ambient control, where CRs were ultimately fulfilled. Available paired observations were used without adjustment for cultivar, season, or stage. Responses to insufficient chilling were expressed as log₂ fold changes between mean concentrations under chilling-deficient and corresponding control conditions. Their magnitude, direction, and consistency were compared across seasons.

## 3. Results

### 3.1 Chilling conditions and associated phenotypic responses

The two experiments provided contrasting chilling conditions for evaluating phytohormone dynamics in apricot floral buds across consecutive seasons. In 2023/2024, chill accumulation at the warm site was 18% lower than at the mild/cold site. In 2024/2025, plastic-film warming increased mean daily maximum temperature by 4.9 °C during the monitoring period (Table S4) and reduced chill accumulation by 35% relative to the ambient control.

Previously published phenotypic data from these experiments are summarized in Table 1 to support interpretation of the hormonal profiles presented here. During the first season, ‘Cebasred’ and ‘Búlida’ showed no substantial flowering disorders at either location, whereas ‘Flopría’ exhibited markedly reduced floral bud viability and cumulative reproductive success at the warm site. Full bloom was delayed, although pistil abortion was lower than at the mild/cold site (Delgado et al., 2026b). During the second season, experimental chill deprivation reduced floral bud viability and cumulative reproductive success in ‘Ladycot’ and increased pistil abortion, despite advancing full bloom (Delgado et al., 2026a). These responses provide the phenotypic context for examining hormonal changes under contrasting chilling conditions.

**Table 1.** Previously published agroclimatic conditions, flowering dates, and reproductive performance of the high-chill apricot cultivars ‘Flopría’ and ‘Ladycot’ under contrasting chilling conditions. ‘Flopría’ was evaluated at mild/cold and warm sites during the 2023/2024 season, whereas ‘Ladycot’ was evaluated under ambient control and plastic-film-induced chill deprivation during the 2024/2025 season. Chill accumulation (CA), expressed in chill portions (CP), was calculated from 1 November to 15 March. Full bloom (F50) was defined as the date on which 50% of flowers were open. Cumulative reproductive success was calculated as the percentage of small fruits relative to the initial number of dormant floral buds. References indicate the source of the data for each season.

| Season | Site/condition | CA (CP) | Cultivar | Viable floral buds (%) | Pistil abortion (%) | Fruit set (%) | Cumulative reproductive success (%) | Full bloom (F50) | Reference |
|---|---|---|---|---|---|---|---|---|---|
| 2023/2024 | Mild/cold | 65.3 | Flopría | 61.2 | 17.0 | 37.6 | 22.3 | 18-Mar | Delgado et al. (2026b) |
|  | Warm | 53.8 | Flopría | 9.0 | 8.0 | 67.2 | 7.1 | 23-Mar |  |
| 2024/2025 | Ambient control | 64.8 | Ladycot | 43.7 | 6.0 | 21.1 | 15 | 25-Mar | Delgado et al. (2026a) |
|  | Chill deprivation | 42.1 | Ladycot | 17.7 | 41.0 | 5.7 | 2.8 | 19-Mar |  |

### 3.2. Phytohormonal variation among cultivars and sampling stages under field conditions

Phytohormonal dynamics during the final stages of CR fulfilment differed among the three apricot cultivars across the two agroclimatic areas. Mean concentrations of several cytokinins, together with SA, JA and IAA-Asp, varied according to cultivar and dormancy progression. Principal component analysis (PCA) showed partial differentiation among cultivars based on their hormonal profiles, with PC1 and PC2 explaining 57.1 and 10.9% of the total variance, respectively (Figure 1). ‘Cebasred’ samples were mainly differentiated along PC1 and generally showed lower PC2 scores. However, this cultivar was sampled only after dormancy release, limiting direct comparisons with ‘Búlida’ and ‘Flopría’, whose sampling periods spanned the dormancy transition. These two cultivars showed partially overlapping hormonal profiles at earlier sampling points but diverged at the final sampling point. A complementary PCA excluding ‘Cebasred’ showed a similar pattern, with PC1 and PC2 explaining 51.1 and 14.1% of the total variance, respectively (Figure S1). Samples from the mild/cold and warm areas showed no consistent separation. Nevertheless, distances between paired samples varied among cultivars and sampling points, suggesting that the influence of agroclimatic conditions depended on genotype and developmental stage.

**Figure 1.**
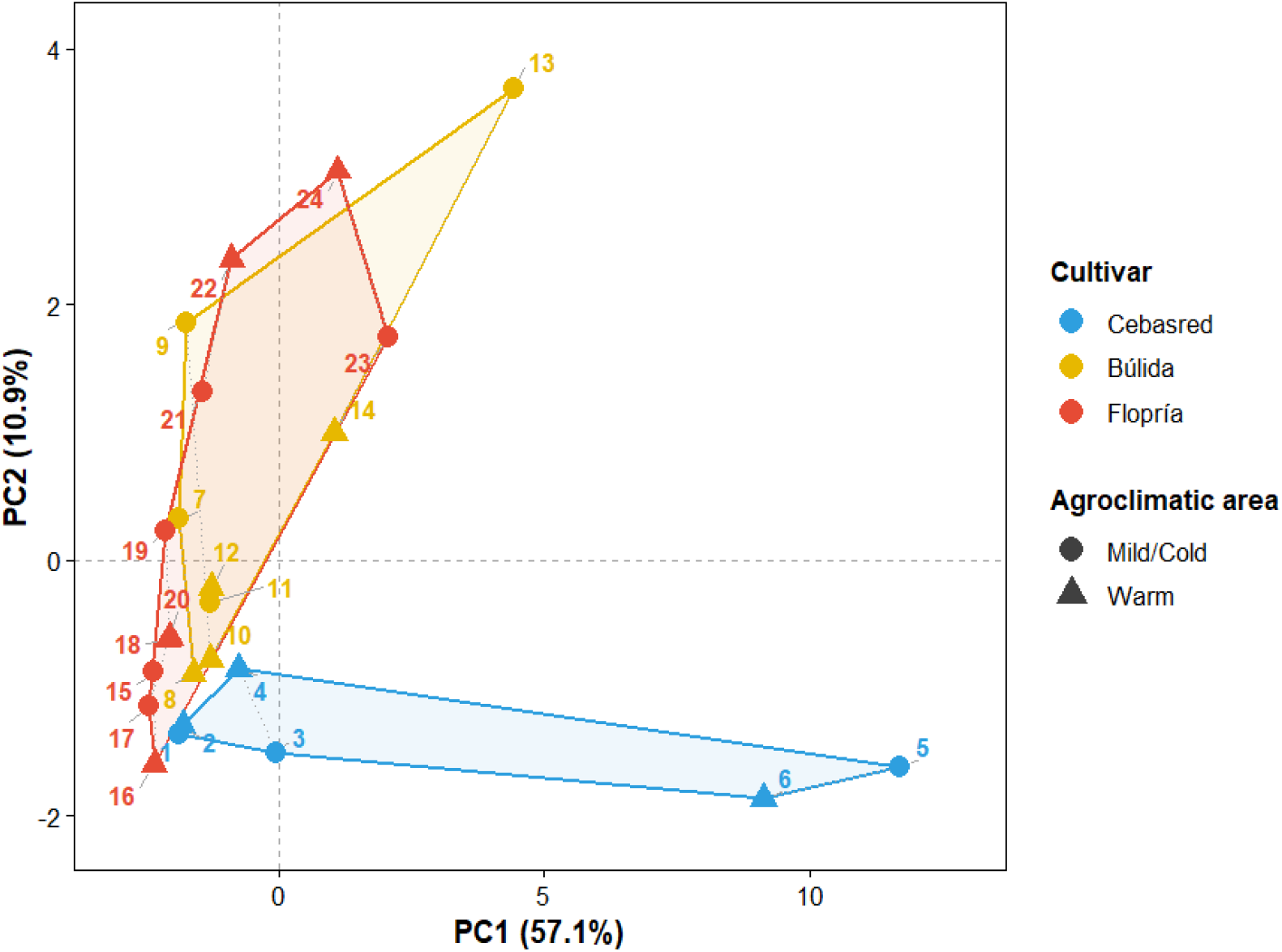
Principal component analysis of phytohormone profiles in apricot floral buds during the late stages of dormancy. Colours identify the three cultivars (‘Cebasred’, ‘Búlida’, and ‘Flopría’), whereas point shapes distinguish the mild/cold and warm agroclimatic areas. Lines connect samples from the two agroclimatic areas corresponding to the same cultivar and sampling point, and convex hulls delimit samples belonging to each cultivar. PC1 and PC2 explained 57.1 and 10.9% of the total variance, respectively. Each point represents the mean phytohormone profile of the independently processed within-tree bud pools. Sample numbers identify unique combinations of cultivar, agroclimatic area, and sampling point, as detailed in Table S5.

### 3.3. Phytohormonal dynamics under contrasting field agroclimatic conditions

Across cultivars, phytohormonal profiles generally followed similar temporal trajectories under the two agroclimatic conditions, although differences in the magnitude of individual responses were observed. In the low-chill cultivar ‘Cebasred’, which had already fulfilled its CR at the beginning of sampling, many phytohormones showed comparable patterns in both areas and increased towards the final assessment at the green tip stage, corresponding to approximately 129–135% CR fulfilment (Figure 2). In ‘Búlida’, phytohormone concentrations generally remained low until endodormancy release and subsequently increased in both areas. However, the magnitude of this increase differed between sites, with N⁶-(Δ²-isopentenyl)adenine (iP), its riboside (iPR) and riboside 5J-monophosphate (iPRMP), trans-zeatin riboside O-glucoside (tZROG), and trans-zeatin riboside 5J-monophosphate (tZRMP) reaching higher concentrations at the mild/cold site than at the warm site during the final sampling stage (Figure 3). JA also showed a more pronounced accumulation peak under mild/cold conditions.

**Figure 2.**
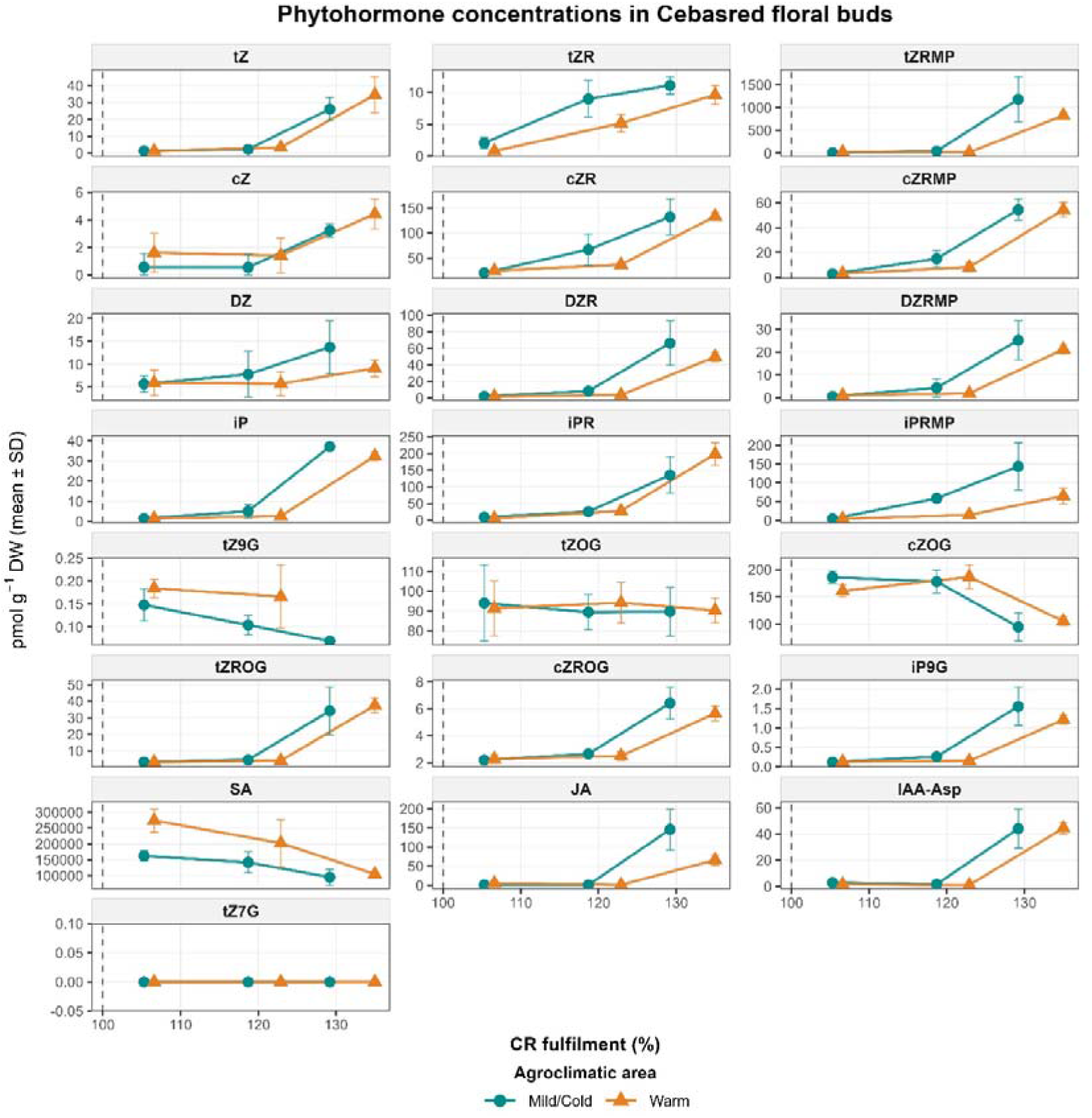
Temporal variation in 22 phytohormone concentrations, comprising 19 cytokinin forms, IAA-Asp, JA, and SA, in floral buds of the low-chill apricot cultivar ‘Cebasred’ under mild/cold and warm agroclimatic conditions during the 2023/2024 season in southeastern Spain. The x-axis represents cultivar- and area-specific percentage of chilling requirement (CR) fulfilment. Points and error bars represent mean concentrations ± SD. The vertical dashed line indicates 100% CR fulfilment, and lines connect consecutive sampling stages within each agroclimatic area. Blue and orange represent the mild/cold and warm areas, respectively. Sampling information is provided in Table S1.

**Figure 3.**
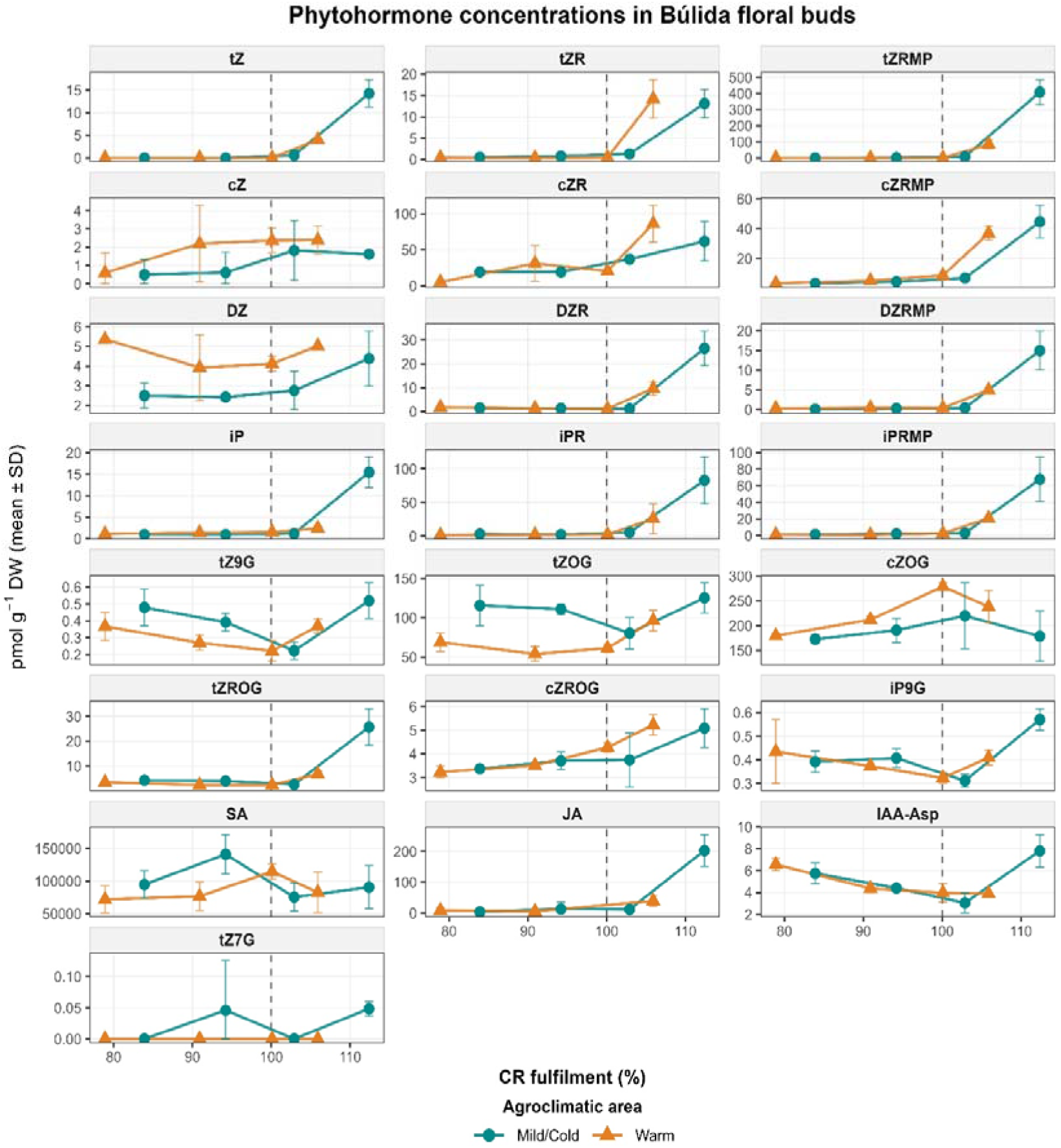
Temporal variation in 22 phytohormone concentrations, comprising 19 cytokinin forms, IAA-Asp, JA, and SA, in floral buds of the medium-chill apricot cultivar ‘Búlida’ under mild/cold and warm agroclimatic conditions during the 2023/2024 season in southeastern Spain. The x-axis represents cultivar- and area-specific percentage of chilling requirement (CR) fulfilment. Points and error bars represent mean concentrations ± SD. The vertical dashed line indicates 100% CR fulfilment, and lines connect consecutive sampling stages within each agroclimatic area. Blue and orange represent the mild/cold and warm areas, respectively. Sampling information is provided in Table S1.

In the high-chill cultivar ‘Flopría’, phytohormone concentrations showed broadly comparable temporal trends in both agroclimatic areas, with relatively low concentrations during the early sampling stages followed by increases around or after CR fulfilment (Figure 4). Differences between agroclimatic areas were mainly associated with the magnitude of the later accumulation peaks rather than the overall trajectories (Figures 2-4). Several cytokinin forms, including iP, iPRMP, DZRMP, tZRMP, and cZR, reached higher final concentrations under mild/cold conditions, whereas other compounds, such as cZ *O*-glucoside (cZOG) and tZROG, showed higher concentrations under warm conditions. JA accumulation around CR fulfilment was also markedly stronger at the mild/cold site. Overall, differences between locations were mainly reflected in the magnitude of late-stage hormone accumulation rather than in the general temporal trajectories.

**Figure 4.**
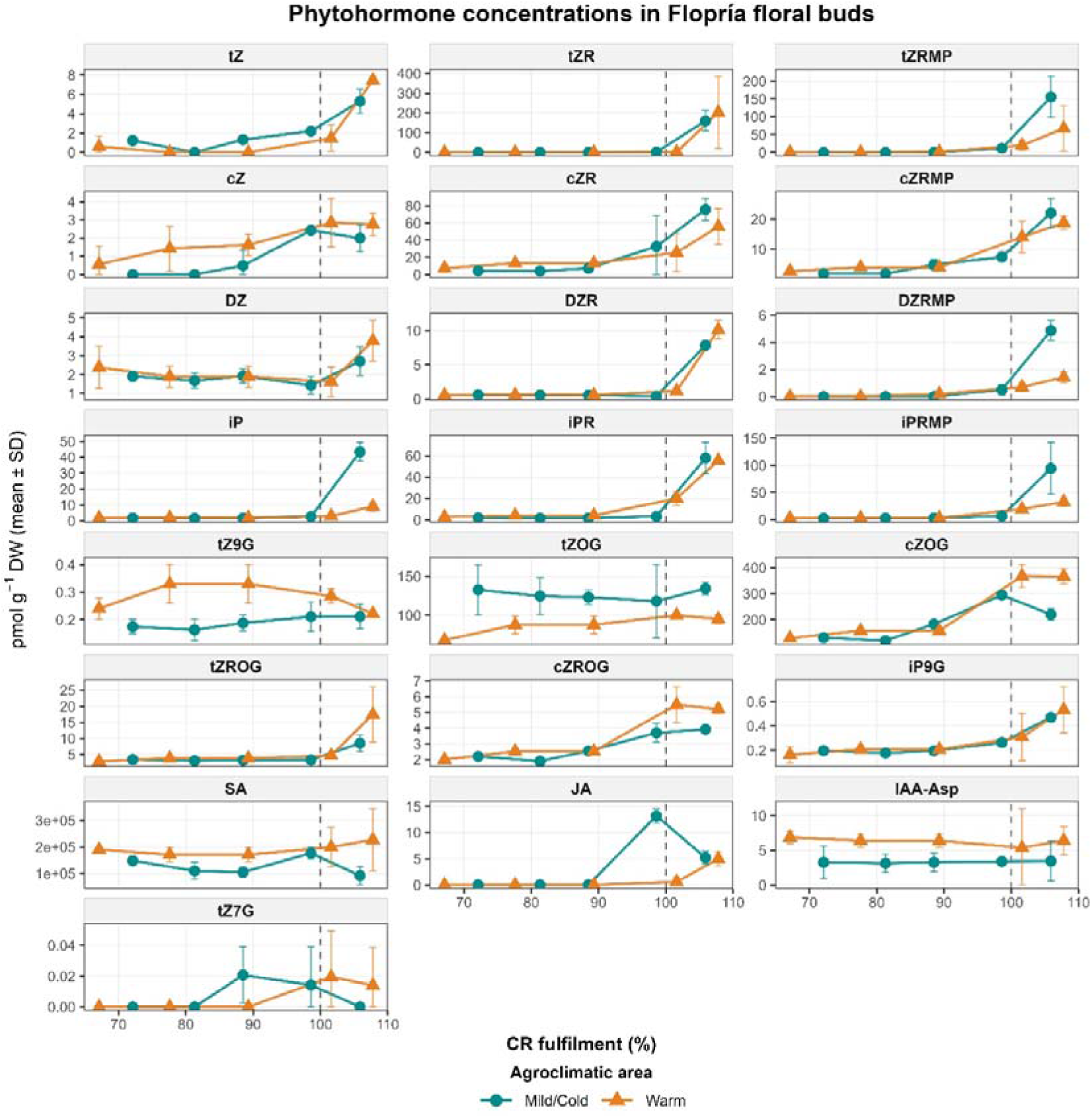
Temporal variation in 22 phytohormone concentrations, comprising 19 cytokinin forms, IAA-Asp, JA, and SA, in floral buds of the high-chill apricot cultivar ‘Flopría’ under mild/cold and warm agroclimatic conditions during the 2023/2024 season in southeastern Spain. The x-axis represents cultivar- and area-specific percentage of chilling requirement (CR) fulfilment. Points and error bars represent mean concentrations ± SD. The vertical dashed line indicates 100% CR fulfilment, and lines connect consecutive sampling stages within each agroclimatic area. Blue and orange represent the mild/cold and warm areas, respectively. Sampling information is provided in Table S1.

The heatmap compared relative phytohormone abundance across cultivars, agroclimatic areas, and sampling stages (Figure 5). In ‘Cebasred’, sampled only after CR fulfilment, several phytohormones reached their highest relative abundance at the final sampling point. In ‘Búlida’, the clearest shift occurred after endodormancy release, between sampling points 2 and 3, with several cytokinins increasing towards the fourth sampling point. In ‘Flopría’, sampling began at approximately 70% CR fulfilment and covered a broader portion of dormancy progression, with several cytokinin forms increasing around or after endodormancy release between sampling points 3 and 4. Differences in cytokinin accumulation between ‘Flopría’ and ‘Búlida’ may suggest cultivar-dependent hormonal responses around endodormancy release. Overall, hormonal patterns varied with cultivar and sampling stage, while agroclimatic conditions modulated the magnitude of specific responses.

**Figure 5.**
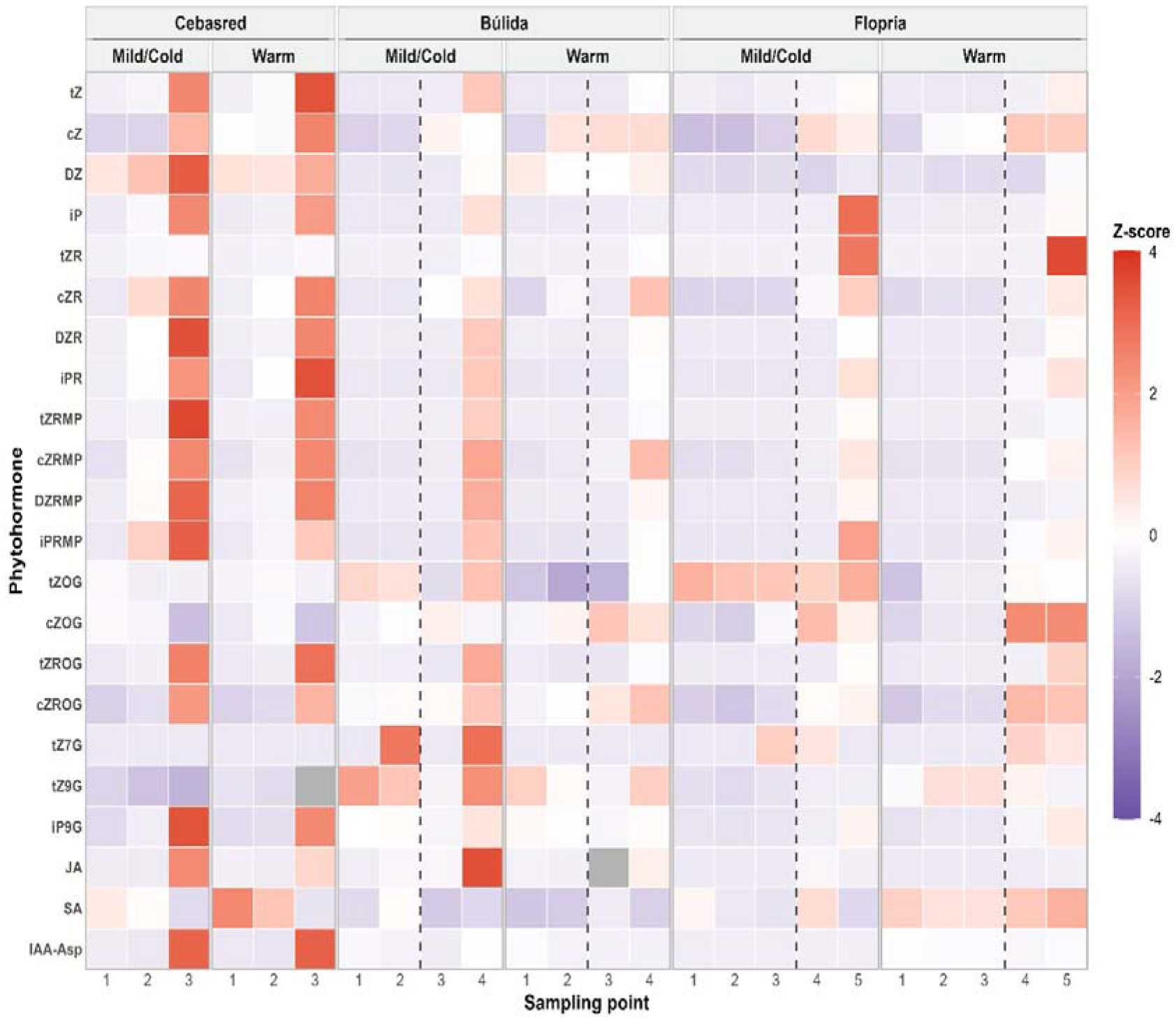
Heatmap of relative abundance of 22 phytohormones in floral buds of low-, medium-, and high-chill apricot cultivars ‘Cebasred’, ‘Búlida’, and ‘Flopría’, respectively, under mild/cold and warm conditions during 2023/2024. Mean concentrations were standardized separately for each phytohormone across all cultivar, area, and sampling-point combinations. Red and purple indicate values above and below the compound-specific mean, respectively; white indicates values near the mean, and grey denotes unavailable measurements. Vertical dashed lines mark endodormancy release between sampling points 2–3 in ‘Búlida’ and 3–4 in ‘Flopría’. No line appears for ‘Cebasred’, sampled only after CR fulfilment. Sampling points represent successive stages of dormancy progression, with cultivar- and location-specific CR fulfilment values provided in Table S1.

### 3.4. Phytohormone responses to experimentally induced severe chill deprivation

#### 3.4.1 Cytokinin content and dynamics during dormancy progression

Most cytokinin forms increased during dormancy progression in both control and chill-deprived trees (Figure 6). However, at the final assessment (126% CR fulfilment in control trees), cZR, dihydrozeatin (DZ), dihydrozeatin riboside (DZR), DZRMP, iPR, and trans-zeatin riboside (tZR) generally reached higher concentrations in control than in chill-deprived buds. This pattern indicates that experimental chill deprivation reduced, delayed, or partially suppressed the late accumulation of several cytokinin forms.

**Figure 6.**
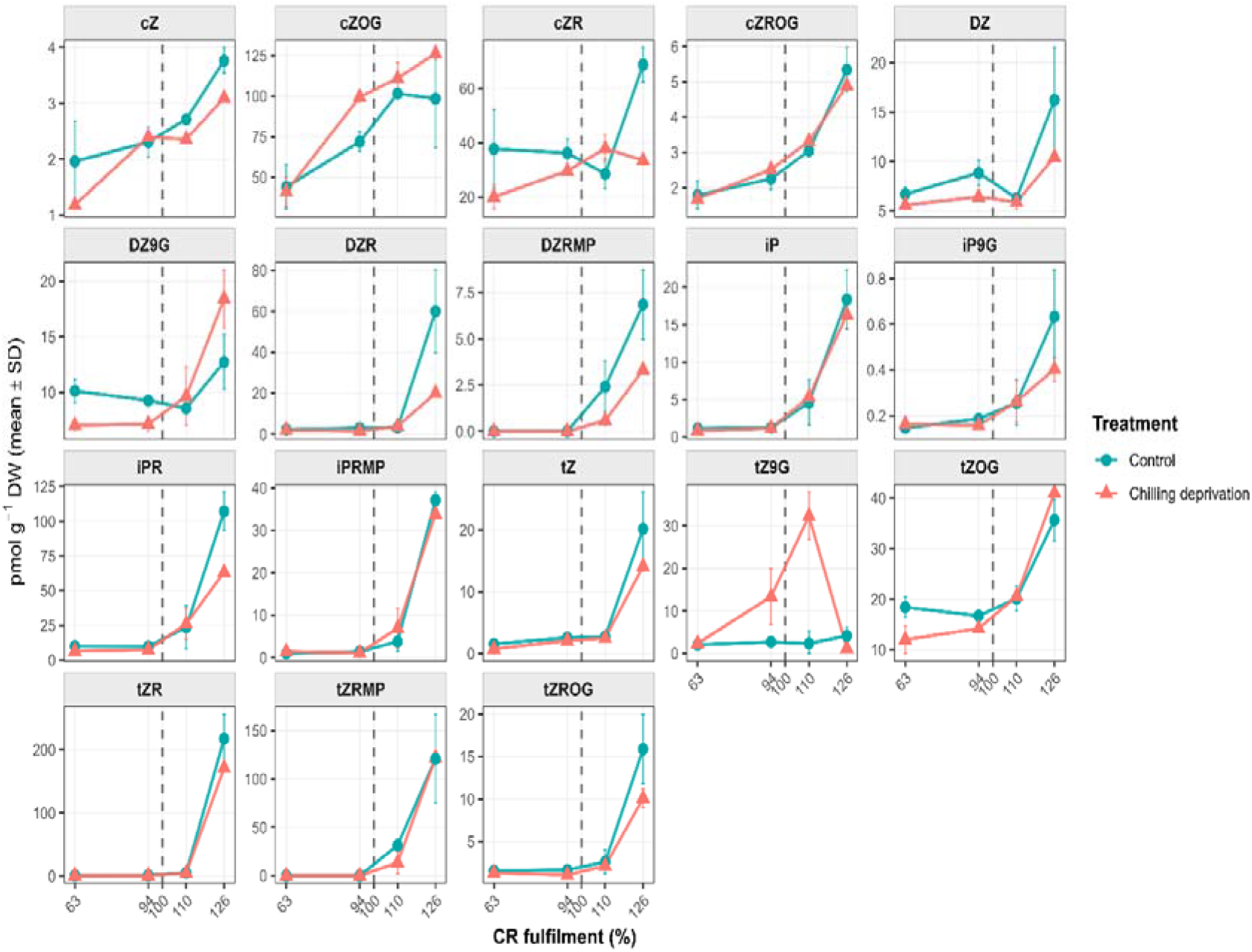
Temporal variation in the concentrations of 18 cytokinin (CK) forms in floral buds of the high-chill apricot cultivar ‘Ladycot’ under ambient control and chill-deprivation conditions. Chill deprivation was induced by prolonged warming under a plastic-film cover. Samples from both treatments were collected on the same calendar dates, corresponding to approximately 63, 94, 110, and 126% of the chilling requirement (CR) estimated for control trees. Blue and red lines represent the ambient control and chill-deprivation treatments, respectively. The vertical dashed line indicates 100% CR fulfilment in the control.

Despite this overall pattern, several cytokinins showed treatment-specific dynamics before the final sampling stage. Notably, trans-zeatin-9-N-glucoside (tZ9G) showed a sharp, transient peak at 110% CR exclusively under chilling deprivation, followed by a rapid decline at 126%, while remaining essentially unchanged in the control (Figure 6). DZ9G concentrations were higher in the control during early sampling stages but only increased in chill-deprived trees at the final stage (126%). The treatments also differed in cZR dynamics: concentrations declined between 94% and 110% in control buds before increasing sharply at 126%, whereas chill-deprived buds showed an increase at 110% followed by a slight decline. By contrast, cZOG generally increased in both treatments, with higher mean concentrations under chill deprivation from 94% onwards.

When only the control treatment was considered, statistically significant changes were identified in iP between 94 and 110% CR fulfilment, the interval encompassing endodormancy release. More extensive changes occurred between 110 and 126%, when 13 cytokinin forms increased significantly (Table S6). These included active bases (iP, tZ, and DZ), ribosides (iPR, tZR, and DZR), nucleotide precursors (iPRMP and tZRMP), and O-glucosides (cZROG, tZOG, and tZROG). Across the broader interval from 94 to 126% CR fulfilment, 15 cytokinin forms changed significantly, further supporting the existence of a coordinated hormonal transition. Furthermore, these findings indicate that extending sampling beyond estimated CR fulfilment improves the detection of coordinated hormonal changes and may facilitate the identification of biologically meaningful indicators of dormancy progression and release.

#### 3.4.2. Dynamics of acidic phytohormones during dormancy progression

During this season, the expanded analytical panel included GA₁, ABA, SA, JA, JA-Ile, IAA, and IAA-Asp. Experimental warming reduced chill accumulation by approximately 35% and modified the temporal profiles of several of these compounds (Figure 7). Overall, hormone concentrations tended to be higher under the colder conditions across most assessment dates. ABA concentrations progressively declined from 63% to 126% CR fulfilment in both treatments, indicating a gradual reduction in dormancy-associated signalling during the transition toward active growth. In contrast, the bioactive gibberellin GA₁ remained low and relatively stable during early stages of CR fulfilment before increasing sharply just after endodormancy release (Figure 7).

**Figure 7.**
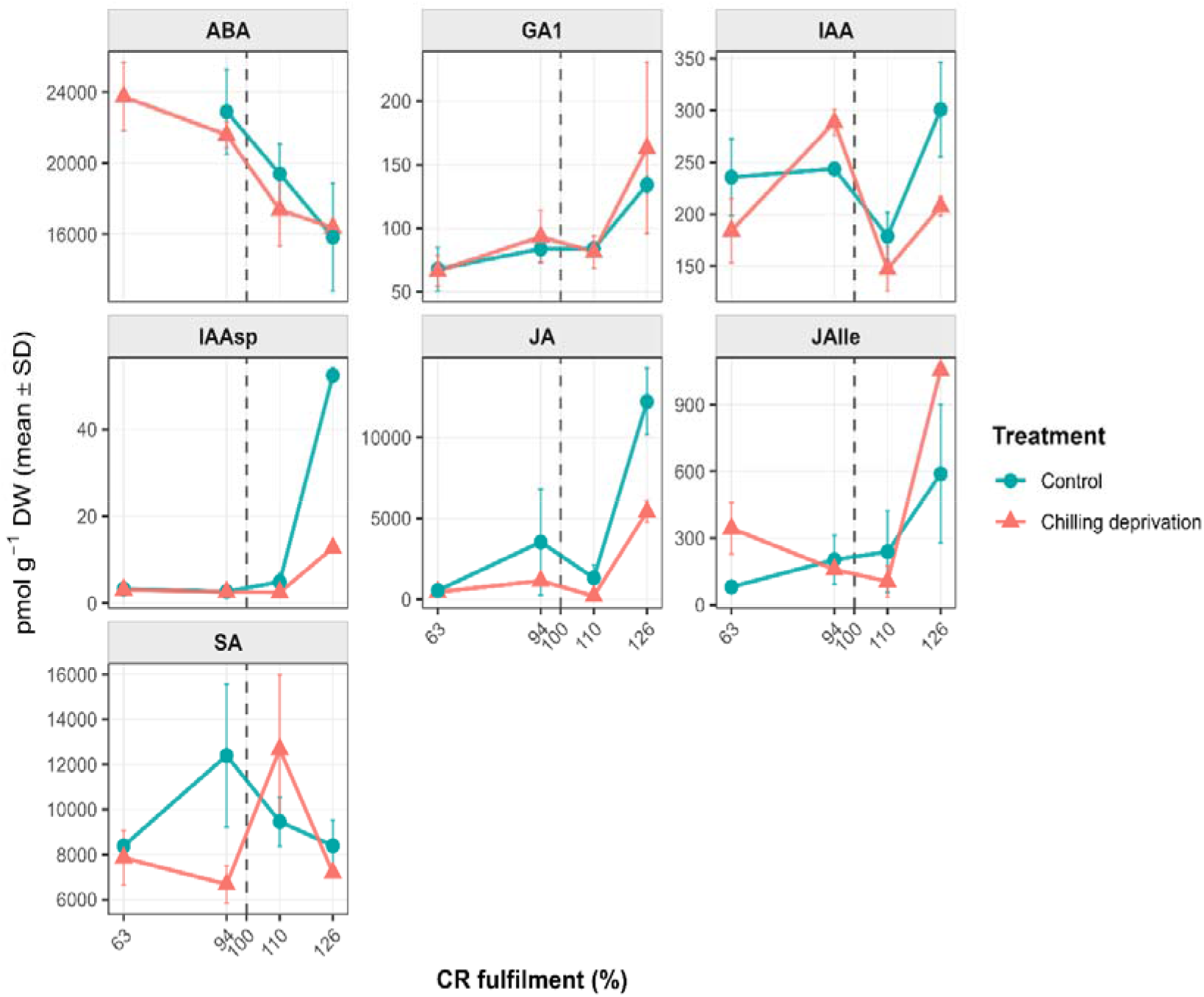
Temporal dynamics of the contents of abscisic acid (ABA), gibberellin A_1_ (GA_1_), indole-3-acetic acid (IAA), indole-3-acetyl aspartate (IAA-Asp), jasmonic acid (JA), jasmonoyl-L-isoleucine (JAlle) and salicylic acid (SA) concentrations (pmol/gDW; Mean ± SD) in ‘Ladycot’ apricot buds. Chill deprivation was induced by prolonged warming under a plastic-film cover. Samples from both treatments were collected on the same calendar dates, corresponding to approximately 63, 94, 110, and 126% of the chilling requirement (CR) estimated for control trees. Blue and red lines represent the ambient control and chill-deprivation treatments, respectively. The vertical dashed line indicates 100% CR fulfilment in the control.

The progressive decrease in ABA and increase in GA₁ concentrations throughout the period examined suggest a coordinated hormonal shift associated with budbreak. Accordingly, the ABA/GA_1_ ratio showed a clear decline during dormancy progression as chilling accumulated, with no statistically significant differences between control and chill-deprived conditions at any stage. Under control conditions, the ABA/GA₁ ratio declined significantly from 231.6 at 94% CR fulfilment to 100.5 at the latest stage (Table S7). A comparable pattern was observed under chill deprivation.

Auxin-related compounds showed dynamic patterns strongly influenced by chilling conditions. IAA levels fluctuated throughout CR fulfilment and increased again at 126% CR fulfilment, although its concentrations were lower under chilling deprivation at this stage. IAA-Asp, a conjugated form of IAA, showed a clearer treatment response, accumulating to higher concentrations at the final stage in control trees. JA also increased markedly at the final stage in control trees, whereas this accumulation was reduced by chill deprivation. In contrast to JA, JA-Ile accumulated more strongly under chill deprivation, reaching its highest concentration at the final sampling stage. Salicylic acid (SA) reached its maximum concentration at 94% CR fulfilment in control trees but peaked at 110% in chill-deprived trees, indicating a temporal shift in its accumulation pattern.

Considering only control trees, which showed normal flowering, potential hormonal indicators of endodormancy release were further assessed. Across the interval from 94 to 110% CR fulfilment, which encompassed endodormancy release, only IAA showed significant changes. Between 110 and 126%, significant changes were detected in IAA, IAA-Asp, GA₁, and JA, whereas the broader transition from 94 to 126% involved significant changes in ABA, JA, and IAA-Asp. These results indicate coordinated changes across several hormonal pathways during the transition from endodormancy release to active bud growth.

### 3.5. Associations among phytohormones during dormancy progression

To further investigate the coordinated regulation of hormonal responses during dormancy transition, Spearman’s rank correlations were calculated using samples from environments in which cultivar-specific CRs were fulfilled: the mild/cold site in the first season and the ambient control in the second season (Figure 8). The resulting matrix revealed predominantly positive correlations among phytohormones. Strong positive associations were observed among several cytokinin forms, including iP, iPR, iPRMP, tZ, tZR, and their conjugates, suggesting closely related changes in their concentrations during dormancy progression. JA-Ile, JA, GA₁, and the auxin-related compounds IAA and IAA-Asp also showed positive associations with several cytokinin forms, although their strength varied among compounds. In contrast, ABA was negatively correlated with most phytohormones, consistent with its role as a dormancy-maintaining signal. Overall, these patterns suggest that dormancy progression under sufficient chilling is associated with coordinated changes in multiple phytohormones, including the accumulation of several growth-related hormones and the concurrent decline in ABA, rather than with independent changes in individual compounds. However, these associations may reflect shared developmental trends and cultivar or seasonal differences rather than direct regulatory interactions.

**Figure 8.**
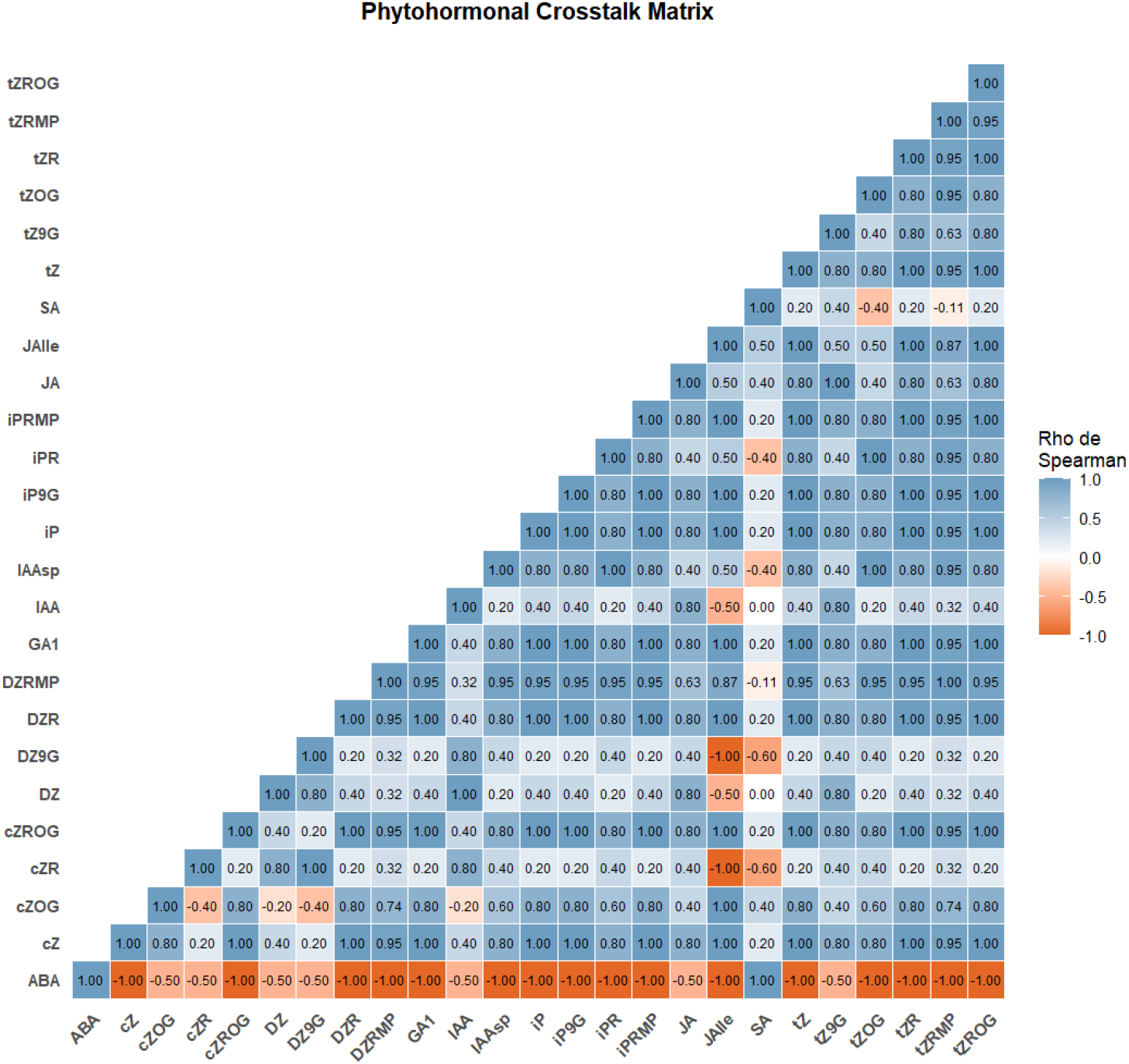
Spearman’s rank correlation matrix of phytohormone concentrations during dormancy progression in apricot floral buds under conditions in which cultivar-specific chilling requirements were fulfilled. The analysis included only samples from trees that ultimately fulfilled their cultivar-specific chilling requirements, including trees at the mild/cold site in the first season and ambient control trees in the second season. Colours indicate the direction and magnitude of the correlations, with orange and blue representing negative and positive associations, respectively. Values within cells are Spearman’s correlation coefficients (ρ).

### 3.6. Candidate phytohormones associated with endodormancy release across experiments

To identify potential indicators of endodormancy release, temporal phytohormone patterns were compared across experiments under conditions in which cultivar-specific CRs were fulfilled, as described in the previous section. Across both experimental seasons, several cytokinins were closely associated with dormancy release under conditions of sufficient chilling. In the first-season orchard experiment, several cytokinin forms, including iP, iPR, iPRMP, tZR, tZRMP, and tZROG, remained at relatively low concentrations during endodormancy and increased markedly following CR fulfilment. In ‘Ladycot’, significant changes between consecutive sampling dates spanning endodormancy release were detected only for iP. Among the compounds examined, iP therefore showed the most consistent increase across cultivars with different CRs and across experimental conditions. Other cytokinin forms showed pronounced responses in the first season but less consistent changes around endodormancy release in the second. These findings identify iP as a promising candidate indicator of endodormancy release under adequate chilling, although its reliability requires further validation.

### 3.7. Phytohormones sensitive to insufficient chilling across experiments

While the previous section focused on hormonal responses under adequate chilling, this analysis examined changes associated with impaired dormancy progression under insufficient chilling. Hormonal profiles under moderate natural and severe experimental chill deficits were compared to identify metabolites responsive to reduced winter chill. Mean phytohormone concentrations across regions differing by approximately 18% in chill accumulation provided limited evidence of widespread hormonal changes under moderate chilling deficiency. Although ‘Flopría’ trees in the warmer region exhibited fewer floral buds and poor cumulative reproductive success (Table 1), these impairments were not accompanied by major changes in the overall phytohormonal profile. Nevertheless, iP, iPRMP, and DZRMP reached higher concentrations at the mild/cold site, where cultivar-specific CRs were fulfilled (Figure 4), whereas cZOG accumulated markedly after endodormancy release at the warm site. JA increased pronouncedly during dormancy progression at the mild/cold site but showed no comparable peak under warmer conditions.

Phytohormones quantified in both experiments were compared to assess whether plastic-film warming reproduced or intensified responses to natural chill deficiency. The more severe experimental deficit (∼35% reduction in chill accumulation) generally reduced hormone concentrations and peak amplitudes relative to controls (Figures 6 and 7). Across both experiments in high-chill cultivars, iP, iPRMP, and DZRMP accumulated more strongly at specific stages following endodormancy release under adequate chilling, while JA increased more when CRs were fulfilled. Conversely, cZOG accumulated under warmer conditions in both seasons, and tZ9G increased under chill deficiency, particularly under severe deprivation. Its transient increase between approximately 63 and 110% CR fulfilment, followed by a return to control levels at the final stage, suggests an association with altered dormancy progression.

Several phytohormones showed consistent mean responses to insufficient chilling across both experiments (Table 2). JA decreased by 50.2 and 56.5% in ‘Flopría’ and ‘Ladycot’, respectively, while tZ decreased by 36.5 and 27.5%. DZRMP also declined in both cultivars, with a larger reduction in ‘Ladycot’ (63.7%) than in ‘Flopría’ (15.5%). Conversely, cZOG increased by 21.6 and 17.3%, respectively. The largest increase occurred for tZ9G, which rose by 52.8% in ‘Flopría’ and 401.0% in ‘Ladycot’. These shared response directions indicate that semi-controlled warming reproduced several hormonal features observed under natural chill deficiency, although their magnitude differed between experiments. These compounds therefore represent candidates for further evaluation as indicators of altered dormancy progression under insufficient chilling.

**Table 2.** Mean phytohormone responses to insufficient chilling in floral buds of two high-chill apricot cultivars evaluated in separate experiments over consecutive seasons. ‘Flopría’ was evaluated under mild/cold and warm field conditions during 2023/2024, whereas ‘Ladycot’ was evaluated under ambient control and plastic-film-induced chill deprivation during 2024/2025. Percentage changes were calculated at each corresponding sampling stage relative to the adequate-chill condition and subsequently averaged across stages. Values are presented for ‘Flopría’ and ‘Ladycot’, respectively. Positive and negative values indicate higher and lower mean concentrations under chill-deficient conditions, respectively.

| Phytohormone class | Candidate indicator | Mean response to insufficient chilling | Magnitude of change (%)<sup>1</sup> | Physiological interpretation |
|---|---|---|---|---|
| Cytokinins | tZ | Consistent decrease | -36.5 / -27.5 | Attenuated accumulation of bioactive cytokinin |
| Cytokinins | DZRMP | Consistent decrease | -15.5 / -63.7 | Reduced accumulation of a cytokinin nucleotide precursor |
| Cytokinins | tZ9G | Strong increase | +52.8 / +401.0 | Increased accumulation of a cytokinin conjugate, With a pronounced transient peak |
| Jasmonates | JA | Strong decrease | −50.2 / −56.5 | Lower mean JA accumulation during dormancy transition |
| Cytokinins | cZOG | Consistent increase | +21.6 / +17.3 | Increased accumulation of a cytokinin conjugate |
| Cytokinins | iPRMP | Consistent increase | +24.7 / +23.8 | Increased cytokinin precursor availability and altered homeostasis |

<sup>1</sup> Values correspond to ‘Flopría’ in 2023/2024 and ‘Ladycot’ in 2024/2025, respectively.

## 4. Discussion

### 4.1. Agroclimatic modulation of hormonal dynamics during dormancy and candidate indicators of endodormancy release

Several phytohormones, including ABA, gibberellins, auxins, cytokinins, and jasmonates, participate in bud dormancy regulation through interconnected pathways whose interactions remain incompletely understood (Chen et al., 2023; Gonzalez Noguer et al., 2023). Accordingly, the contrasting phenological and reproductive responses among apricot cultivars, and the variation in estimated CRs across environments, probably reflect the integration of temperature cues with genotype-dependent hormonal regulation rather than the action of individual pathways. By combining field and semi-controlled experiments, this study characterized phytohormone dynamics from late endodormancy to active bud growth under different levels of chill accumulation. Most phytohormones followed broadly similar temporal trajectories across environments, while reduced chilling primarily modified the magnitude and timing of their accumulation. Comparable observations in *Prunus* suggest that climatic variation can modify the intensity of dormancy-related responses while largely maintaining their overall seasonal progression (Hernandez et al., 2021; Herrera et al., 2026).

Regarding cultivar-specific variation in hormone concentrations, cytokinin dynamics varied among cultivars and developmental stages. Sapkota et al. (2021) reported a positive association between cytokinin levels and flowering time in apple. In partial agreement, the high-chill, later-flowering ‘Flopría’ generally showed higher cytokinin concentrations around endodormancy release than the medium-chill ‘Búlida’.

Several cytokinin forms remained at low concentrations during endodormancy and increased after CR fulfilment, consistent with their roles in cell division, meristem activity, and sink establishment during growth resumption (Sakakibara, 2006). Cytokinins may also interact antagonistically with gibberellins during floral transition by promoting GA degradation and repressing GA signalling, as reported in apple (Li et al., 2019). Although limited cytokinin variation during dormancy has been reported in sweet cherry (Götz and Chmielewski, 2023), our results consistently showed increases in iP, iPR, iPRMP, tZR, tZRMP, and DZR after dormancy release across cultivars and experimental seasons. Partly consistent with our observations, Ionescu et al. (2017) reported increases in tZR, DZ, and DZR during hydrogen cyanamide-induced bud break in sweet cherry, while tZ and tZR increased during natural bud break in poplar (Hewett and Wareing, 1973). Collectively, these findings support the involvement of specific cytokinins in the transition from dormancy to active growth in woody perennials. However, the magnitude and consistency of these changes differed among cytokinin forms, highlighting the importance of evaluating individual compounds rather than the total cytokinin pool. Among the cytokinins evaluated in our study, iP showed the most consistent response across cultivars and experiments, supporting its potential as an indicator of endodormancy release in apricot.

Among acidic phytohormones, JA and JA-Ile increased towards the final sampling stage but responded differently to chill deprivation: JA accumulation was reduced, whereas JA-Ile concentrations increased. Previous studies suggest that jasmonates may contribute to bud break in deciduous woody perennials through the regulation of cell division, vascular reconnection, and tissue differentiation processes (Ionescu et al., 2017; Liu and Sherif, 2019). Similarly, IAA and IAA-Asp increased after dormancy release, consistent with the activation of auxin-related pathways during bud growth resumption in grapevine (Noriega and Pérez, 2017). Hiejima et al. (2025) suggested that SA depletion was associated with ABA reduction during dormancy release in Japanese cedar. Unlike the progressive decline in ABA, SA showed a transient peak whose timing shifted under chill deprivation, suggesting distinct temporal responses of these hormones. The increase in JA after CR fulfilment and its positive associations with cytokinins, auxins, and GA₁ suggest its involvement in hormonal reprogramming during dormancy release and growth reactivation.

In contrast to the rapid activation of most cytokinins and jasmonates, ABA and GA₁ exhibited complementary but temporally distinct dynamics. ABA declined progressively under both conditions, supporting the association between declining ABA levels and dormancy release in agreement with studies in sweet cherry (Chmielewski et al., 2017), peach and apricot (Guillamón Guillamón et al., 2024; Guillamón et al., 2025). Conversely, GA₁ remained low during chilling accumulation and increased sharply once CR was fulfilled. Thus, the decline in the ABA/GA₁ ratio during dormancy release mainly reflected decreasing ABA rather than GA₁ accumulation, consistent with the progressive attenuation of dormancy-maintaining signals reported in *Prunus* spp. (Hernandez et al., 2021; Zhang et al., 2024). GA₁ therefore appears more closely associated with growth activation than with endodormancy release, as reported in sweet cherry (Götz et al., 2023) and Japanese apricot (Zhang et al., 2018). Collectively, the temporal patterns of selected cytokinins, jasmonates, auxins, and GA₁, together with their negative associations with ABA, suggest interrelated hormonal changes during this transition. Selected cytokinins may therefore serve as physiological indicators of chilling fulfilment, although validation across genotypes, seasons, and environments remains necessary. Multi-year assessments are particularly important because warm winter periods may alter response magnitude while preserving overall temporal pattern.

### 4.2. Hormonal alterations during dormancy progression under insufficient chilling

Several studies have associated reproductive failure with insufficient winter chill (Delgado et al., 2026a; Petri and Leite, 2004). However, the molecular and hormonal mechanisms linking chilling deprivation with flowering disorders remain poorly understood (Tominaga et al., 2022). By evaluating two high-chill apricot cultivars with similar CRs under contrasting chill accumulation conditions, ranging from sufficient chilling that supported normal reproductive development to insufficient chilling associated with reproductive failure, this study provides insight into how chill availability modifies hormonal responses during dormancy progression and subsequent bud growth.

Although the main hormonal trajectories were broadly conserved across environments, several hormones were sensitive to chill availability and coincided with reduced flowering intensity and cumulative reproductive success under warmer conditions (Delgado et al., 2026b, 2026a). Among cytokinins, iP, iPRMP, and DZRMP accumulated at higher concentrations under colder conditions, particularly around dormancy release, whereas cZOG showed a marked post-dormancy increase under insufficient chill conditions. These contrasting patterns indicate that chill deprivation modifies the balance among cytokinin forms rather than uniformly reducing their concentrations. JA showed the strongest differential response, with a clear increase around dormancy release occurring only under colder conditions. Together, these results suggest that insufficient chill selectively altered jasmonate- and cytokinin-related responses, potentially affecting the hormonal coordination required for subsequent floral development. The attenuated cytokinin and jasmonate responses under insufficient chilling may reflect impaired meristematic reactivation and floral development, potentially contributing to bud abortion and reduced fruit set under warm winter conditions (Kumar et al., 2017; Tominaga et al., 2022).

The progressive decline in ABA across treatments suggests that it was not strictly dependent on CR fulfilment. The broadly conserved decline in the ABA/GA₁ ratio indicates that chill deprivation did not prevent this hormonal transition but may have altered its timing and coordination. Similarly, Ito et al. (2021) observed declining ABA concentrations in Japanese pear under different chill accumulation regimes and proposed that ABA may be more closely associated with dormancy depth and flowering synchrony than with acting as a binary regulator of growth resumption. The delayed SA peak under chill deprivation further suggests that insufficient chilling altered the timing rather than the occurrence of this hormonal response.

Auxin dynamics were also sensitive to temperature. Under sufficient chilling, IAA increased during late developmental stages, coinciding with JA accumulation, whereas both responses were attenuated under warmer conditions, suggesting weaker coordination among growth-related hormonal pathways. Given the roles of auxins and jasmonates in tissue differentiation, vascular reconnection, and potentially bud growth activation (Wasternack and Hause, 2013), their reduced accumulation under limited chilling conditions may negatively affect bud viability. Nevertheless, the broadly similar hormonal trajectories across treatments suggest that some dormancy-associated developmental processes continued under chill-deficient conditions, despite reduced reproductive performance. The contrasting responses of specific cytokinin forms, particularly the mean decreases in tZ and DZRMP and increases in cZOG and tZ9G, together with the pronounced reduction in JA, identify candidate hormonal responses associated with insufficient chilling and impaired reproductive failure. Further research should clarify their roles in flowering failure under warmer winters. However, the treatment combined reduced chilling with higher pre-bloom temperatures, preventing attribution of the observed responses solely to chill deficiency.

### 4.3. Implications for apricot breeding and climate adaptation strategies

The need to select cultivars adapted to specific agroclimatic conditions, together with the continuous release of new cultivars, highlights the importance of identifying reliable biological indicators associated with endodormancy release. Such indicators could improve the estimation of cultivar-specific CRs and support breeding programmes (Herrera et al. 2022; Delgado et al. 2024). To our knowledge, this study represents the first comprehensive evaluation of a broad range of phytohormones within a narrow temporal window before and after dormancy release across contrasting climatic conditions in *Prunus* spp. Assuming that the bud-forcing method accurately reflects the physiological timing of endodormancy release (Delgado et al., 2026b, 2026a), the significant hormonal changes detected between approximately 94% and 110% CR fulfilment narrowed the range of potential candidates. Only the iP increase was reproducible across cultivars and seasons, supporting its potential as a physiological indicator. Extending the sampling window slightly beyond complete CR fulfilment improved the detection of statistically significant hormonal transitions, although extending this window could reduce the precision of cultivar-specific CR estimates (Delgado et al. 2024). Although hormonal analysis requires specialized equipment and costs more than bud-forcing assays, its potentially shorter turnaround could complement the approximately 10-day forcing period and inform timely applications of dormancy-breaking compounds or plant growth regulators (PGRs).

Beyond their potential use as indicators, attenuated cytokinin and jasmonate responses under insufficient chilling provide a rationale for investigating PGRs as tools to mitigate chill-deficiency symptoms. In particular, our results suggest that cytokinin-mediated signalling, potentially interacting with jasmonate pathways, contributes to a successful dormancy transition. Exogenous application of cytokinins during late dormancy stages may help restore hormonal balance under insufficient chilling conditions, potentially improving budbreak uniformity and flowering intensity. Jasmonate-based treatments may also represent a promising approach by modifying both growth and stress responses. Their effectiveness and optimal timing nevertheless require careful evaluation given the context-dependent functions of jasmonates regulating plant growth (Wasternack and Hause, 2013).

Direct application of endogenous cytokinin forms such as iP, iPRMP, or DZRMP, is uncommon under field conditions. However, synthetic analogues such as 6-benzyladenine (BA) and forchlorfenuron (CPPU) are widely used in horticulture to stimulate cell division, promote bud outgrowth, and improve fruit set (Davies, 2004). For instance, BA has reduced flower abortion in lupin (Atkins and Pigeaire, 1993) and promoted branching in apple (Wertheim and Estabrooks, 1994). In apple rootstocks, 6-BA also promoted bud dormancy release alongside changes in cytokinin and ABA levels (Asghar et al., 2026). Similarly, combinations of gibberellins (e.g., GA₃) and cytokinins have been associated with substantial yield increases in blackberry (Galindo-Reyes et al., 2016), demonstrating the potential of combined PGR applications. Nevertheless, this study did not test exogenous hormones, and the observed associations do not establish that treatments would restore normal dormancy release or reproductive performance in apricot. Orchard trials could compare compounds, concentrations, and application timings and assess their effects on budbreak synchrony, floral viability, and fruit set.

## 5. Conclusions

This study characterizes phytohormone dynamics during the transition from endodormancy to active growth in apricot floral buds under contrasting agroclimatic conditions. Across cultivars, seasons, and environments, hormonal profiles followed broadly consistent temporal trajectories, with pronounced changes after CR fulfilment. ABA concentrations and the ABA/GA₁ ratio declined during dormancy progression, while increases in selected cytokinins, JA, auxins, and GA₁ accompanied growth resumption. However, declining ABA alone may not distinguish successful dormancy progression from incomplete or poorly coordinated development under insufficient chilling. Chill deprivation did not prevent growth resumption entirely but attenuated or delayed several hormonal responses. Under severe deprivation, these changes coincided with reduced bud viability and reproductive performance, indicating that conserved hormonal trajectories can coexist with impaired development. iP showed the most reproducible increase across cultivars and experiments, supporting its candidacy as an indicator of endodormancy release. JA may provide complementary information on growth activation and chill deficiency. Consistent mean decreases in tZ and DZRMP and increases in cZOG and tZ9G also identify candidate responses for characterising insufficient chilling. These findings could help refine cultivar-specific CR assessment and support cultivar selection for warmer winters. Broader genetic and environmental validation, together with functional analyses of hormone biosynthesis and signalling, remains necessary before practical application in breeding and climate-adaptation strategies.

## Acknowledgements

The authors gratefully acknowledge Jesús López Alcolea (CEBAS-CSIC) for his assistance with orchard selection, Diego Moreno (CEBAS-CSIC) for his support in sample preprocessing, and the private fruit companies El Ciruelo® and FERMONDUC S.A. for providing access to their commercial orchards. The authors also thank R-COMS_CSRS_YC3191, YC0967 for supporting the plant hormone analysis platform.

## Authors’ contributions

AD, DR, and HY designed the research. AD and JAS collected the plant material. AD and TFH prepared the samples. HS developed the analytical platform for hormone identification. MK performed the UPLC–qMS/MS metabolomic analyses. RSP and JAE supervised the data analysis and contributed to the interpretation of the results. AD wrote the manuscript. All authors read and approved the final version of the manuscript.

